# WTFgenes: What's The Function of these genes? Static sites for model-based gene set analysis

**DOI:** 10.1101/114785

**Authors:** Christopher J. Mungall, Ian H. Holmes

## Abstract

A common technique for interpreting experimentally-identified lists of genes is to look for enrichment of genes associated to particular ontology terms. The most common technique uses the hypergeometric distribution; more recently, a model-based approach was proposed. These approaches must typically be run using downloaded software, or on a server. We develop a collapsed likelihood for model-based gene set analysis and present WTFgenes, an implementation of both hypergeometric and model-based approaches, that can be published as a static site with computation run in JavaScript on the user's web browser client. Apart from hosting files, zero server resources are required: the site can (for example) be served directly from Amazon S3 or GitHub Pages. A C++11 implementation yielding identical results runs roughly twice as fast as the JavaScript version. WTFgenes is available from https://github.com/evoldoers/wtfgenes under the BSD3 license. A demonstration for the Gene Ontology is usable at https://evoldoers.github.io/wtfgo. Contact: Ian Holmes.

## Introduction

Term Enrichment Analysis (TEA) is a common technique for finding functional patterns, specifically over-represented ontology terms, in a set of experimentally identified genes (Boyle *et al.*, 2004). The most common approach, which we refer to as *Frequentist TEA,* is a one-tailed Fisher's Exact Test (based on the hypergeometric distribution, which models the number of term-associations if the gene set was chosen by chance), with a suitable correction for multiple hypothesis testing. Frequentist TEA has been implemented many times on various platforms (Robinson *et al.*, 2002; Khatri *et al.*, 2002; Zeeberg *et al.*, 2003; Boyle *et al.*, 2004; Bauer *et al.*, 2008; Jiao *et al.*, 2012; Mi *et al.*, 2013; Chen *et al.*, 2013).

A model-based alternative to Frequentist TEA, which more directly addresses some of the multiple testing issues (for example, by modeling the ways that an observed gene list can be broken down into complementary gene sets), is *Bayesian TEA*. In contrast to Frequentist TEA, which just rejects a null hypothesis that genes are chosen by chance, the Bayesian TEA explicitly models the alternative hypothesis that the gene set was generated from a few random ontology terms. This approach was introduced by Lu *et al.* (2008) and further developed by Bauer *et al.* (2010), who implemented model-based testing in Java and R (Bauer *et al.*, 2011). However, the model-based approach remains significantly less well-explored than frequentist approaches.

The graphical model underpinning Bayesian TEA is sketched in Figure 1. For each of the *m* terms there is a boolean random variable *T_j_* (“term *j* is activated”). For each of the *n* genes there is a directly-observed boolean random variable *O_i_* (“gene *i* is observed in the gene set”), and one deterministic boolean variable *H_i_* (“gene *i* is activated”) defined by *H_i_* = 1 − Π_*j*∈*G_i_*_ (1 − *T_j_*) where *G_i_* is the set of terms associated with gene *i* (including directly annotated terms, as well as ancestral terms implied by transitive closure of the directly annotated terms). The probability parameters are *π* (term activation), *α* (false positive) and *β* (false negative), and the respective hyperparameters are **p** = (*p*_0_, *p*_1_), **a** = (*a*_0_, *a*_1_) and **b** = (*b*_0_, *b*_1_). The model is

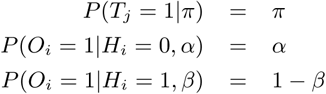

with *π* ~ Beta(**p**), *α* ~ Beta(**a**) and *β* ~ Beta(**b**). The model of Bauer *et al.* (2010) is similar but used an *ad hoc* discretized prior for *π*, *α* and *β*.

**Figure 1:**
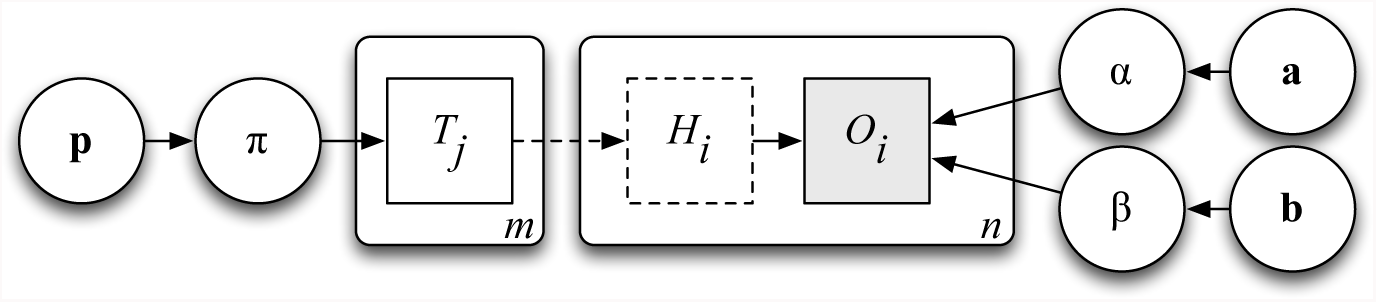
Model-based explanation of observed genes (*O_i_*) using ontology terms (*T_j_*), following Bauer *et al.* (2010). Other variables and hyperparameters are defined in the text. Circular nodes indicate continuous-valued variables or hyperparameters; square nodes indicate discrete-valued (boolean) variables. Dashed lines indicate deterministic relationships; shaded nodes indicate observations. Plates (rounded rectangles) indicate replicated subgraph structures.

Most Bayesian and Frequentist TEA implementations are designed for desktop use. Several Frequentist TEA implementations are designed for the web, such as DAVID-WS (Jiao *et al.*, 2012) and Enrichr (Chen *et al.*, 2013) which has a rich dynamic web front-end. However, web-facing Frequentist TEA implementations generally require a server-hosted back end that executes code. Further, there are no web-based Bayesian TEA implementations.

## Results

In developing our Bayesian TEA sampler, we introduce a collapsed version of the model in Figure 1 by integrating out the probability parameters. Let 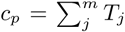 count the number of activated terms, 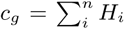 the activated genes, 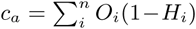 the false positives and 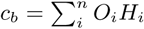 the false negatives. Then

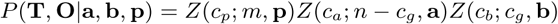

where

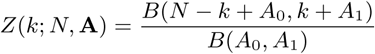

is the beta-Bernoulli distribution for *k* ordered successes in *N* trials with hyperparameters **A** = (*A_0_*, *A*_1_), using the beta function

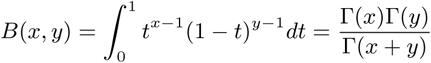

Integrating out probability parameters improves sampling efficiency and allows for higher-dimensional models where, for example, we observe multiple gene sets and give each term its own probability *π_j_* or each gene its own error rates (*α_i_*, *β_i_*). Our implementation by default uses uninformative priors with hyperparameters **a** = **b** = **p** = (1, 1) but this can be overridden by the user.

The MCMC sampler uses a Metropolis-Hastings kernel (Gilks *et al.*, 1996). Each proposed move perturbs some subset of the term variables. The moves include *flip*, where a single term is toggled; *step*, where any activated term and any one of its unactivated ancestors or descendants are toggled; *jump*, where any activated term and any unactivated term are toggled; and *randomize*, where all term variables are uniformly randomized. The relative rates of these moves can be set by the user.

The sampler of Bauer *et al.* (2010) implemented only the *flip* move. To test the relative efficacy of the newly-introduced moves we measured the autocorrelation of the term variables for one of the GO Project's test sets, containing 17 *S.cerevisiae* mating genes^1^. The results, shown in Figure 2, led us to set the MCMC defaults such that the *flip*, *step*, and *jump* moves are equiprobable, while *randomize* is disabled.

**Figure 2:**
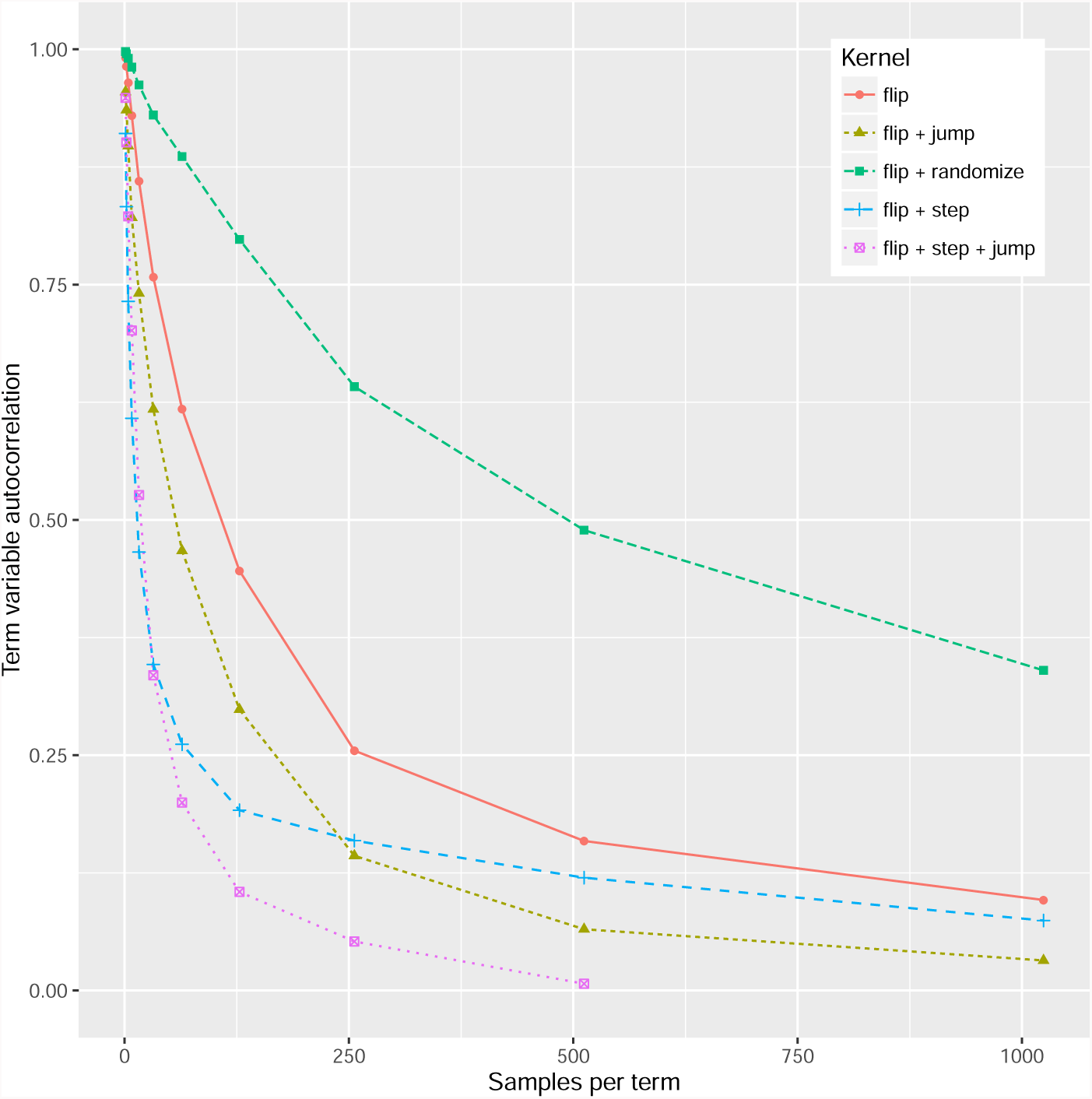
Autocorrelation of term variables, as a function of the number of MCMC samples, for several MCMC kernels on a set of 17 *S. cerevisiae* mating genes. A rapidly-decaying curve indicates an efficiently-mixing kernel. The kernel incorporating *flip*, *step* and *jump* moves (defined in the text) mixes most efficiently.

We have implemented both Frequentist TEA (with Bonferroni correction) and Bayesian TEA (as described above), in both C++11 and JavaScript. The JavaScript version can be run as a command-line tool using node, or via a web interface in a browser, and includes extensive unit tests. The two implementations use the same random number generator and yield numerically identical results. The C++ version is about twice as fast: a benchmark of Bayesian TEA on a late-2014 iMac (4GHz Intel Core i7), using the abovementioned 17 yeast mating genes and the relevant subset of 518 GO terms, run for 1,000 samples per term, took 37.6 seconds of user time for the C++ implementation and 79.8 seconds in JavaScript.

By contrast, the Frequentist TEA approach is almost instant. However, its weaker statistical power is apparent from Figure 3, which compares the recall *vs* specificity of Bayesian and Frequentist methods on simulated datasets. For values of N from 1 to 4, we sampled N terms from the *S.cerevisiae* subset of the Gene Ontology, and generated a corresponding set of yeast genes with false positive rate 0.1% and false negative rate 1%. The MCMC sampler was run for 100 iterations per term, and this experiment was repeated 100 times. The model-based approach has vastly superior recall to the Fisher exact test, and the difference grows with the number of terms.

**Figure 3:**
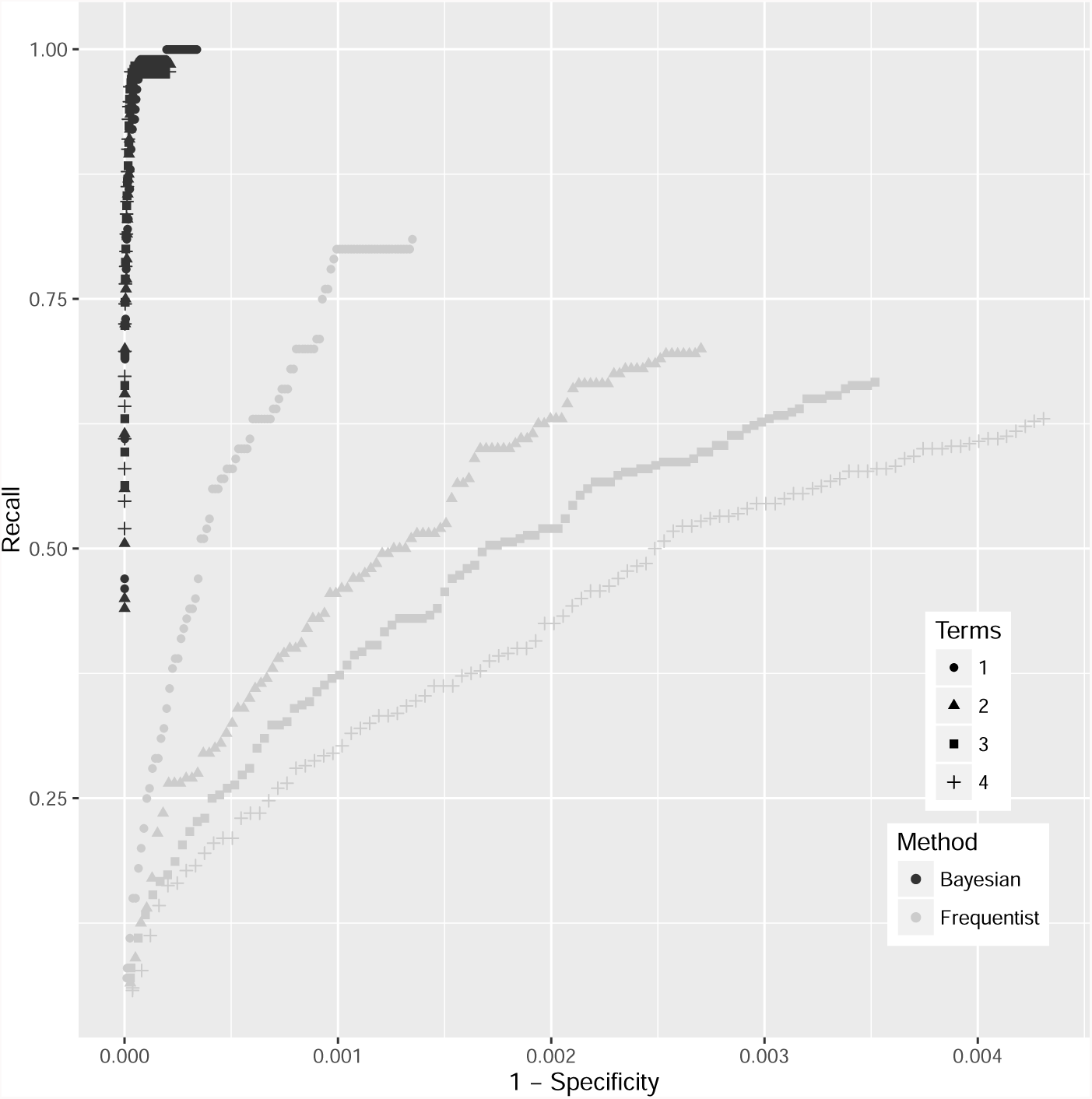
ROC curves for Frequentist and Bayesian TEA. The axes are scaled per term. There are 5,919 ontology terms annotated to *S.cerevisiae* genes, so (for example) a false discovery rate of 0.001 corresponds to about 6 falsely reported terms.

Our JavaScript software, when used as a web application, offers a “quick report” view using Frequentist TEA. For the slower-running but more powerful Bayesian TEA, the software plots the log-likelihood during an MCMC sampling run, for visual feedback. The repository includes setup scripts allowing the tool to be deployed as a “static site”, i.e. consisting only of static files (HTML, CSS, JSON, and JavaScript) that can be hosted via a minimal web server with no need for dynamic code execution. This has considerable advantages: static web hosting is generally much cheaper, and far more secure, than running server-hosted web applications.

An example wtfgenes static site, configured for the GO-basic ontology and GO-annotated genomes from the Gene Ontology website, can be found at https://evoldoers.github.io/wtfgo.

## Discussion

JavaScript genome browsers such as JBrowse (Buels *et al.*, 2016) represent a broader web trend of producing static sites where possible, for reasons of security and performance. We have implemented such a static site generator for ontological term enrichment analysis of gene sets that offers both Bayesian and frequentist tests. In contrast with existing web services for Frequentist TEA, such as DAVID-WS or Enrichr, it requires no server resources and allows comparison of Bayesian and Frequentist approaches.

Model-based TEA is versatile: it can readily be extended to allow for datasets that are structured temporally (Hejblum *et al.*, 2015), spatially (Lin *et al.*, 2015), or by genomic region (McLean *et al.*, 2010); to use domain-specific biological knowledge (Szczurek and Beerenwinkel, 2014); or to incorporate additional lines of evidence such as quantitative data (Kalaitzis and Lawrence, 2011). We hope our development of a collapsed likelihood, and evaluation of different MCMC kernels, will assist these efforts.

Coincidentally, Fisher's Exact Test—which we call Frequentist TEA— was originally motivated by a blind tea-tasting challenge (Fisher, 1935).

## Funding

IHH was partially supported by NHGRI grant HG004483. CJM was partially supported by Office of the Director R24-OD011883 and by the Director, Office of Science, Office of Basic Energy Sciences, of the US Department of Energy under Contract No. DE-AC02-05CH11231.

## Footnotes

1 Gene IDs: STE2, STE3, STE5, GPA1, SST2, STE11, STE50, STE20, STE4, STE18, FUS3, KSS1, PTP2, MSG5, DIG1, DIG2, STE12.

